# Strategies to improve scFvs as crystallization chaperones suggested by analysis of a complex with the human PHD-bromodomain SP140

**DOI:** 10.1101/767376

**Authors:** Michael Fairhead, Charlotta Preger, Edvard Wigren, Claire Strain-Damerell, Elena Ossipova, Mingda Ye, Mpho Makola, Nicola A. Burgess-Brown, Helena Persson, Frank von Delft, Susanne Gräslund

## Abstract

Antibody fragments have great potential as crystallization chaperones for structural biology due to their ability to either stabilise targets, trap certain conformations and/or promote crystal packing. Here we present an example of using a single-chain variable fragment (scFv) to determine the previously unsolved structure of the multidomain protein SP140. This nuclear leukocyte-specific protein contains domains related to chromatin-mediated gene expression and has been implicated in various disease states. The structure of two of the domains (PHD-bromodomain) was solved by crystallizing them as a complex with a scFv generated by phage display technology. SP140 maintains a similar overall fold to previous PHD-bromodomains and the scFv CDR loops predominately interact with the PHD, while the framework regions of the scFv makes numerous interactions with the bromodomain. Analysis of our and other complex structures suggest various protein engineering strategies that might be employed to improve the usefulness of scFvs as crystallization chaperones.

## Introduction

Despite extensive efforts in construct design (1, 2) and screening of chemical space (3), many targets routinely fail to yield the protein crystals necessary for structural studies *via* x-ray diffraction. This failure may be due to poor protein stability (4), multiple conformational states leading to sample heterogeneity or it even being the wrong time of year (5)! Antibody fragments have proven useful in overcoming some of these issues by acting as crystallization chaperones (6). They can stabilise and/or trap specific conformations of proteins as well as directly mediate crystal contacts. Mimetic binders based on non-immunoglobulin scaffolds have also been engineered and proven useful in this application (7).

Examples of target:binder complexes in the PDB are numerous and steadily growing, with several different binder types having been used, such as Fabs, scFvs, nanobodies and Darpins (see Table 1). Currently there is no clear consensus which binder format is best and most likely this will vary from target to target. For structural studies binders should have a stable and rigid scaffold and regions (such as loops) that tolerate high sequence variability that generates specificity for a target. Ideally binders should also be produced in a way that gives rise to a homogenous preparation (*i.e.* without glycosylations and other modifications).

**Table 1.** Number of binder and binder:complex structures* in the PDB (21st June 2019)

| Binder class | Number of structures<sup>1</sup> | Number of complexes<sup>2</sup> |
| --- | --- | --- |
| Affibody | 13 | 10 |
| Affimer | 9 | 2 |
| Darpin | 141 | 70 |
| Fab | 2502 | 1646 |
| Nanobody | 293 | 207 |
| scFv (+Fv)<sup>4</sup> | 165 (255) | 118 (158) |

\*The search outcomes were not specifically curated beyond the basic search criteria. <sup>1</sup>The type of “binder class” was used to search the full text of the mmCIF file deposited with the PDB. <sup>2</sup>The type of “binder class” & the term “complex” was used to search the PDB. Searches were made using the advanced search page on the PDB ( http://www.rcsb.org/pdb/search/advSearch.do).<sup>4</sup> Although we searched for scFv complexes in the PDB several examples of Fv complexes were found. These have been included in the analysis to increase the number of examples studied. Observations with these binder types should be translatable to the scFv arrangement.

A single chain fusion of the variable heavy (V_H_) and light (V_L_) domains (connected via a peptide linker) of an immunoglobulin (Ig) (8, 9) can fulfil many of the above requirements. The complementary determining regions (CDR) on the surface of the V_H_ and V_L_ domains together generate the antigen-binding site, with the potential of providing high specificity and affinity to most proteins and other types of antigens (10–13). ScFvs with new specificities can be readily generated and selected via techniques such as phage display (14) and if necessary they can be further engineered for increased performance, for example by increasing its stability (15) or affinity (16, 17). We chose the epigenetic target, SP140, as a case study to help develop a pipeline that has the ambition of routinely generating scFvs that recognize natively folded proteins and which can be used as crystallization chaperones.

SP140 is a nuclear leukocyte specific protein containing chromatin reader modules (Figure 1A). So far only the NMR structure of one of these modules (plant homeodomain, PHD) (18) and the PHD-bromodomain x-ray crystal structure of the related SP100C (19) and SP100A (4PTB) proteins have been reported. Although the Structural Genomics Consortium (SGC) has historically been successful in solving the structures of many other bromodomains and related chromatin reader modules (20) the structure of SP140 had remained unsolved. Using an scFv, generated by phage display, we solved the structure of the PHD-Bromodomain of SP140 (6G8R). As well as generating a novel structure, we analysed this complex and those reported by others to investigate some of the features of scFvs, which may aid crystallization and points the way to possible improvements to this scaffold as a crystallization chaperone.

**Figure 1:**
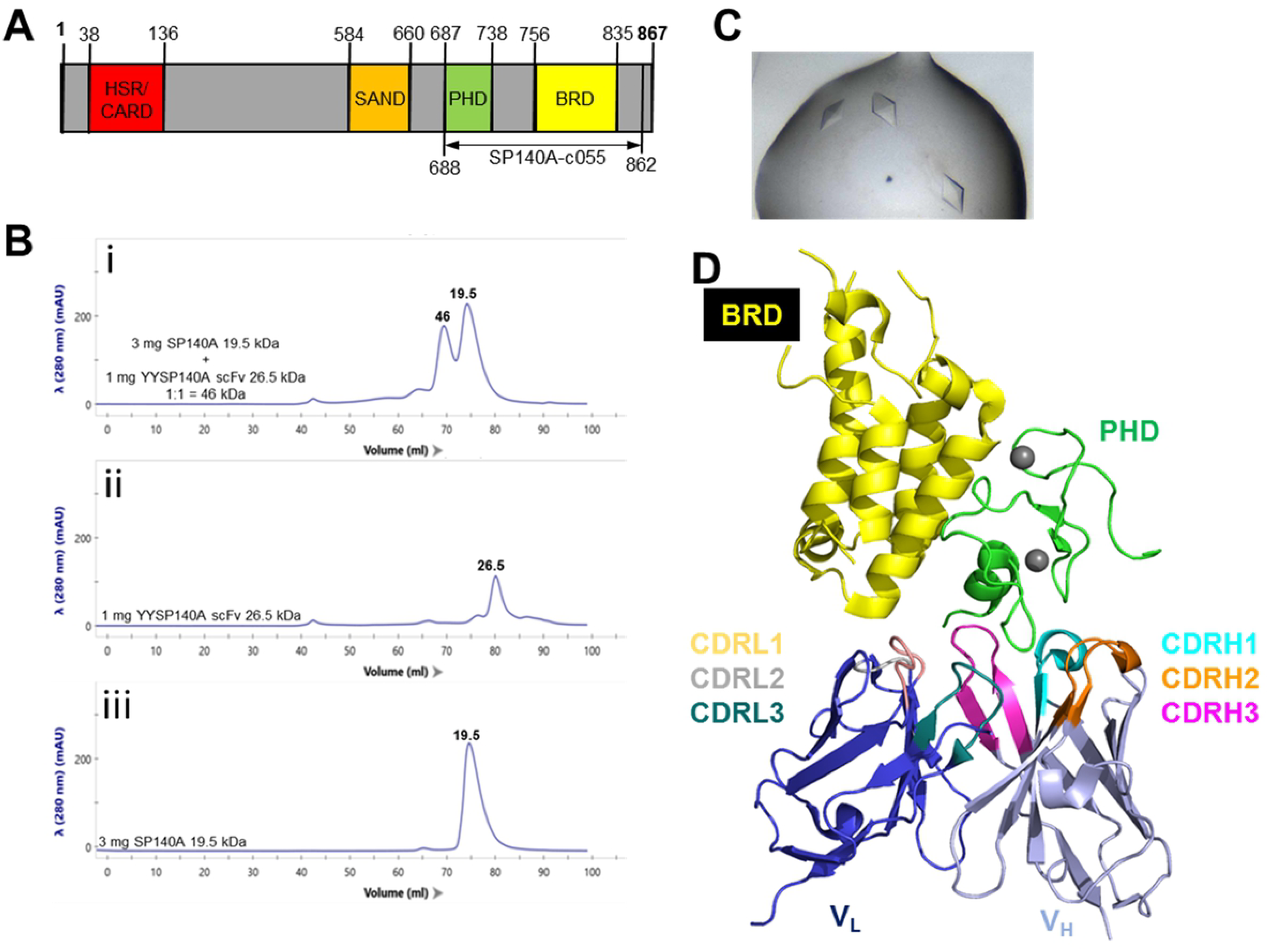
Isolation, crystallization and structure of the scFv:SP140 complex. (A) Overview of domains in human SP140. (B) Size exclusion chromatograms of (i) scFv:SP140A mix, (ii) scFv and (iii) SP140 respectively. (C) Crystals of the scFv:SP140 complex obtained in 1.2 M sodium malonate, 0.5 % jeffamine ED-2003, 0.1 M HEPES pH 7. (D) Cartoon representation (Pymol) of 6G8R with the PHD-bromodomain of SP140, the scFv V_H_ and V_L_ and their respective CDR loops coloured as indicated.

## Results and Discussion

### scFv generation, expression and purification

ScFv binders against the PHD-bromodomain of SP140 (Supplementary Method 1) were generated by phage display technology using a human synthetic library denoted SciLifeLib with biotinylated recombinant SP140 protein as target protein. The top candidate, G-SP140-8, with a measured affinity of approximately 6 nM (Supplementary Figure S1) was selected and used in co-crystallization experiments with SP140.

Although scFvs are useful binder scaffolds, we have found that production in *Escherichia coli* routinely gives low yields of less than 1 mg/L of cells for some if not most constructs. As crystallography can require several milligrams of protein, we evaluated different strategies to improve yield and purity. The most successful strategy was to produce the G-SP140-8 scFv as an MBP fusion and express it in the oxidising cytoplasm of the mutant *E. coli* strain Shuffle (NEB) (21). Isolation of the MBP-scFv fusion with Ni-Sepharose and Protein A (22) resins yielded highly pure protein. Subsequent MBP removal by His_6_-tagged TEV protease followed by reverse chromatography with Ni-Sepharose and amylose resins (to remove MBP, uncleaved protein and his tagged TEV protease) routinely generated more than 5 mg of purified scFv per litre.

### Complex isolation and crystallization

After mixing an excess of SP140 with scFv, the resulting complex was readily isolated via size exclusion chromatography (SEC) (Figure 1B i-iii). A clear shift in molecular weight can be observed for the complex as compared to the individual components, (Figure 1B i-iii). Curiously the isolated scFv was always found to run lower than its predicted MW (Figure 1B ii). This may indicate some dissociation of the two domains during the run (V_H_ and V_L_), *i.e.* the protein overall behaves more like a long linear molecule rather than a single larger globular protein. After isolation by SEC, the complex of scFv:SP140 was put directly into crystallization trials and crystals with a bipyramid morphology were found in many conditions after 1-2 days at room temperature. Unfortunately, most of these crystals were found to diffract to a low resolution (3-5 Å). Only crystals grown in a mixture of malonate/jeffamine (Figure 1C.) were found to diffract to a resolution of 2.7 Å and allow the generation of a structure with reasonable refinement statistics, Table 2.

**Table 2.** Data collection and refinement statistics.

|  | **6G8R** |
| --- | --- |
| **Wavelength** | 0.96861 Å |
| **Resolution range** | 51.32 - 2.74 (2.84 - 2.74) Å |
| **Space group** | P 61 2 2 |
| **Unit cell** | 89.2x89.2x343.35 Å 90x90x120 ° |
| **Total reflections** | 421847 (42464) |
| **Unique reflections** | 22245 (2164) |
| **Multiplicity** | 19.0 (19.6) |
| **Completeness (%)** | 99.46 (99.36) |
| **Mean I/sigma(I)** | 11.89 (1.02) |
| **Wilson B-factor** | 72.26 |
| **R-merge** | 0.2313 (3.242) |
| **R-meas** | 0.2376 (3.327) |
| **R-pim** | 0.05388 (0.7413) |
| **CC1/2** | 0.998 (0.652) |
| **CC\*** | 1 (0.889) |
| **Reflections used in refinement** | 22226 (2160) |
| **Reflections used for R-free** | 1139 (107) |
| **R-work** | 0.2243 (0.3616) |
| **R-free** | 0.2665 (0.3482) |
| **CC(work)** | 0.918 (0.756) |
| **CC(free)** | 0.905 (0.679) |
| **Number of non-hydrogen atoms** | 3057 |
| **macromolecules** | 3033 |
| **ligands** | 10 |
| **solvent** | 14 |
| **Protein residues** | 393 |
| **RMS(bonds)** | 0.003 Å |
| **RMS(angles)** | 0.9 ° |
| **Ramachandran favoured (%)** | 93.67 |
| **Ramachandran allowed (%)** | 6.33 |
| **Ramachandran outliers (%)** | 0 |
| **Rotamer outliers (%)** | 0.93 |
| **Clashscore** | 2.54 |
| **Average B-factor** | 74.49 |
| **macromolecules** | 74.55 |
| **ligands** | 72.99 |
| **solvent** | 61.43 |

2 Statistics for the highest-resolution shell are shown in parentheses.

### Structure of SP140 PHD:Bromodomain

The packing observed in the crystal structure consists of a crystallographic dimer of SP140 (Chain B) surrounded by four copies of the scFv (Chain A) (Supplementary Figure S5). Specifically, the structure of the SP140 PHD:Bromodomain is very similar to that observed for the related SP100 (4PTB) and SP100C (5FB1) proteins with less than a 0.7-0.8 Å RMSD between the structures (Supplementary Figure S6). This small difference is perhaps unsurprising giving the relatively high sequence identity of 61 % (Supplementary sequence S3).

It has previously been demonstrated that the isolated PHD cannot bind the substrate histone tail peptide (18) in the absence of the bromodomain (19). This suggests that the bromodomain is necessary to stabilise the conformation of the PHD and that this stabilization is required to allow binding (19). The structure of the SP140 PHD:Bromodomain presented here confirms this observation as the PHD adopts a conformation similar to that observed in two related PHD:Bromodomain structures, the substrate peptide bound SP100C (5FB1), (19) and the apo SP100 (4PTB). This is in contrast to the isolated PHD NMR structures which have distinct cis or trans conformations (18), neither of which resemble the SP140 PHD:Bromodomain structure (Supplementary Figure S5).

### Interaction of scFv with SP140

As would be expected for an antibody-based binder, residues from the CDR-loops on the surface of scFv form numerous interactions with SP140 (Figure 1D and Figure 2A), predominantly through the PHD. PISA (23) analysis shows binding of the scFv to SP140 results in an extensive interface covering a buried surface area of 842 Å^2^ and of the six CDR-loops, CDR-H3 (395 Å^2^) dominates the interface followed by CDR-H2 (118 Å^2^) and CDR-L3 (86 Å^2^) (Supplementary Figure S7). None of the CDR loops occupy the peptide-binding site of the PHD domain (Figure 2D) suggesting that the scFv would not inhibit protein activity. More unexpected is the importance of the interactions of residues on the side of the scFv opposite to the CDR-loops, which participate in crystal contacts with neighbouring copies of SP140 (Figure 2B). Crystal packing is also observed between neighbouring scFv molecules (Figure 2C).

**Figure 2:**
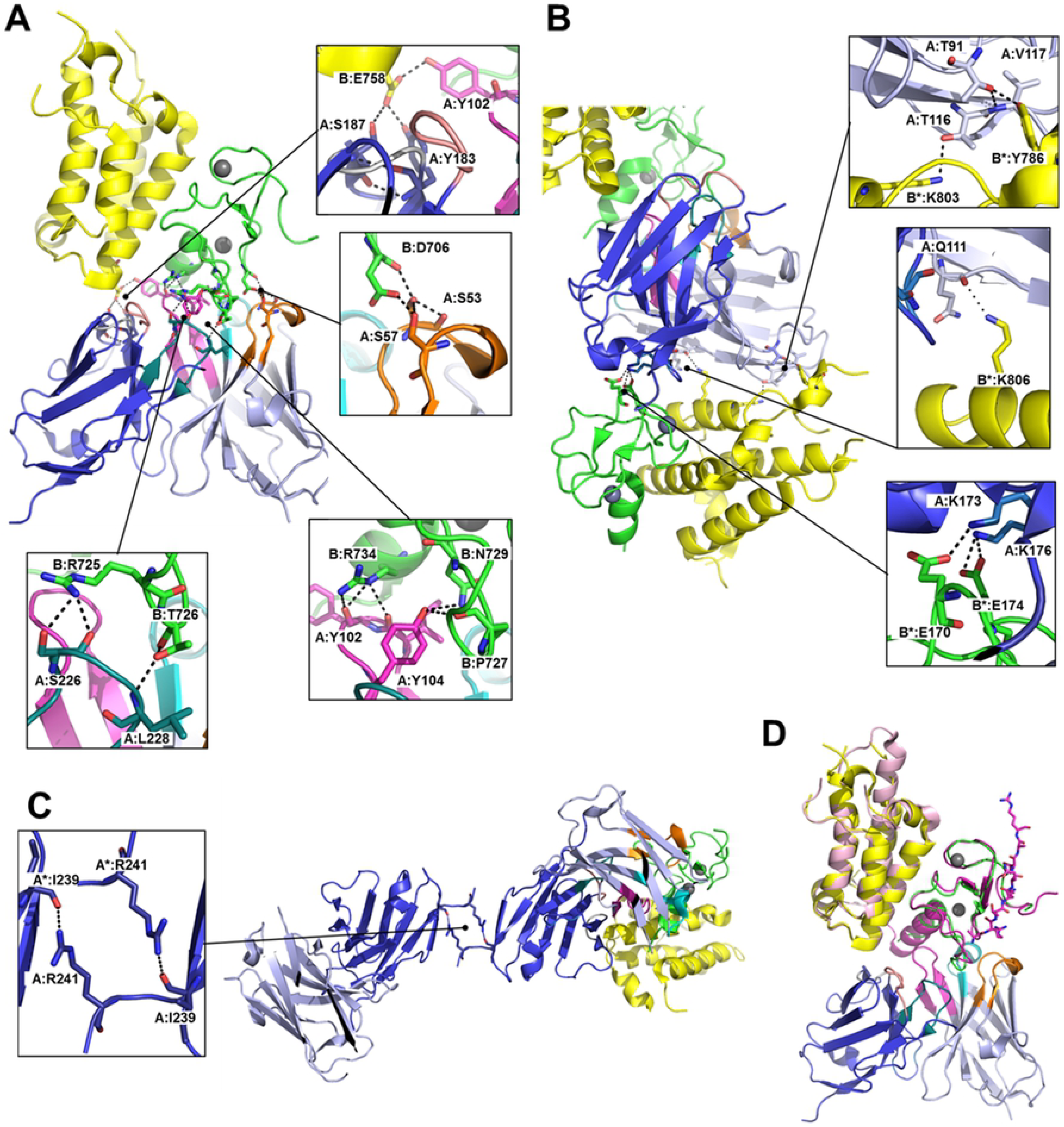
Binding and packing features of the scFv:SP140 complex. Cartoon representation (Pymol) of (A) CDR loop interactions of the scFv with surface residues of SP140. (B) Interactions of the backside of the scFv with a symmetry copy of SP140 (B*). (C) Interactions of the C-terminal tail of the scFv with a symmetry copy (A*). (D) overlay of 6G8R coloured as in Figure 1D with peptide bound SP100C structure (5FB1) with bound peptide in stick representation (hot pink) and the PHD and bromodomains coloured hot pink and pale pink respectively.

The interaction of the scFv with both the bromodomain and PHD and its probable stabilising effect may be important given the above noted fact that the PHD in isolation appears to be non-functional (18). This might explain our failure to crystallize the isolated protein, despite success with many similar classes of protein (20).

### scFvs as crystallization chaperones

Of the binder:complex structures reported in the PDB (Table 1) sixty-one of these contain an scFv or Fv in complex with another protein, the remainder being small molecules, peptides, apo or anomalies (see Table S1 for details). As can be seen from Table S2, these binders have been used in isolation and with other antibody fragments such as Fabs to solve a wide variety of proteins from small toxins (24) to large virus assemblies (25, 26) and a range of proteins in between, demonstrating their utility as crystallization chaperones. While studies have shown that the crystallization properties of the scFv can be modified to produce new space groups, until recently this has been limited to scFvs binding small molecules (27) or peptides (28). However, a recent example demonstrates the potential benefits of engineering the scFv (as well as a Fab) for improved crystal packing, allowing a structural study of small molecules bound to the gp120 subunit of the HIV-1 envelope trimer (29). We reasoned that an analysis of our and others protein:scFv/Fv complexes would yield suggestions on how to generate scFvs with improved crystallization properties.

Table S3 shows a summary of information on the scFv/Fv and their amino acid sequences in these complexes. It can be seen from Table S3 that different investigators have arranged the V_L_ and V_H_ domains from different species in a variety of orders (*i.e.* V_L_ or V_H_ can be first in the scFv) with different linkers and tags. It can also be seen that in some cases the V_L_ and V_H_ domains were sufficiently stable as a complex to not require a linker (Fv rather than scFv).

A simple analysis of the complexes (Table S3) shows that many regions of the scFv constructs are not visible in the final crystal structure; these include long C-terminal purification tags (*e.g.* His_6_, Avi tags). Usually such flexible tags are removed from targets when setting up crystal trials as they are typically disordered and rarely visible in the final structure. The increased entropic cost associated with this disorder reduces the likelihood of crystallization (30). It would be best if the N- and C-termini of the scFv were trimmed to their approximate Fv boundaries.

Another observation is that the V_H_-V_L_ linker is rarely visible in structures (Table S3). If this linker could be redesigned to be more rigid for the above reasons of entropic cost, this may increase crystallization success rates. One approach may be to simply increase the serine content, which has been reported to increase stiffness (31). Alternatively a computational approach to the design could be taken (32). Care has to be taken as linker modifications can have unintended adverse effects such as reducing affinity, stability and oligomerization (33, 34), any of which which may be disadvantageous for crystallization and target binding.

In Table S4 an analysis of the regions commonly found in Fv crystal contacts (not involving the primary Fv:target complex) shows that many crystal packing arrangements of Fv molecules are possible. These multiple possible packing arrangements may partly explain the usefulness of these binders as crystallization chaperones.

A summary of the various Fv crystal contacts and an alignment of the different V_L_ and V_H_ domains (IMGT numbering done using the ANARCI program (35)) is shown in Tables S5 and S6. Although there are regions which appear to be quite frequently observed at crystal contacts (*e.g.* around residues 5-15 in V_L_), a limitation is that the number of examples of complexes analysed is relatively small (61 in total) and that the Fv domains are from a number of species (human, mouse, rhesus, rabbit & rat), meaning a high degree of sequence variability. Even with these limitations it is perhaps surprisingly how frequently the same regions are found in crystal packing points with other Fv (Tables S5 and S6). Targeting of these regions with mutagenesis is likely a good strategy for improving scFv as crystallization chaperones.

Analysis of the alignments in Tables S5 and S6 also shows that the Fv:Fv packing frequently involve a glutamine, lysine or glutamate. These residues are associated with high entropic cost and mutating these to alanine or a residue with lower entropy (arginine, aspartate or asparagine) has been shown in other proteins to be a successful strategy to improve crystallization success (36). Such a surface entropy reduction (SER) strategy has been done to improve the crystallization properties of an EE epitope (EYMPME) binding scFv (28), although without a target protein bound. Similarly a SER approach has been used on a Fab scaffold to improve crystallization when in complex with RNA (37).

### Conclusions

The structure of SP140 in complex with scFv presented here shows that the PHD adopts the expected fold as observed in previous PHD:Bromodomain structures such as SP100 (4PTB) and SP100C (5FB1) The success of this crystallization attempt is likely a result of stabilization of the PHD due to the presence of interactions with both the bromodomain and the scFv. Analysis of the scFv in this and previously reported structures suggests that the utility of this class of binders as routine crystallization chaperones would be greatly improved through certain modifications, summarized in Figure 3.

**Figure 3:**
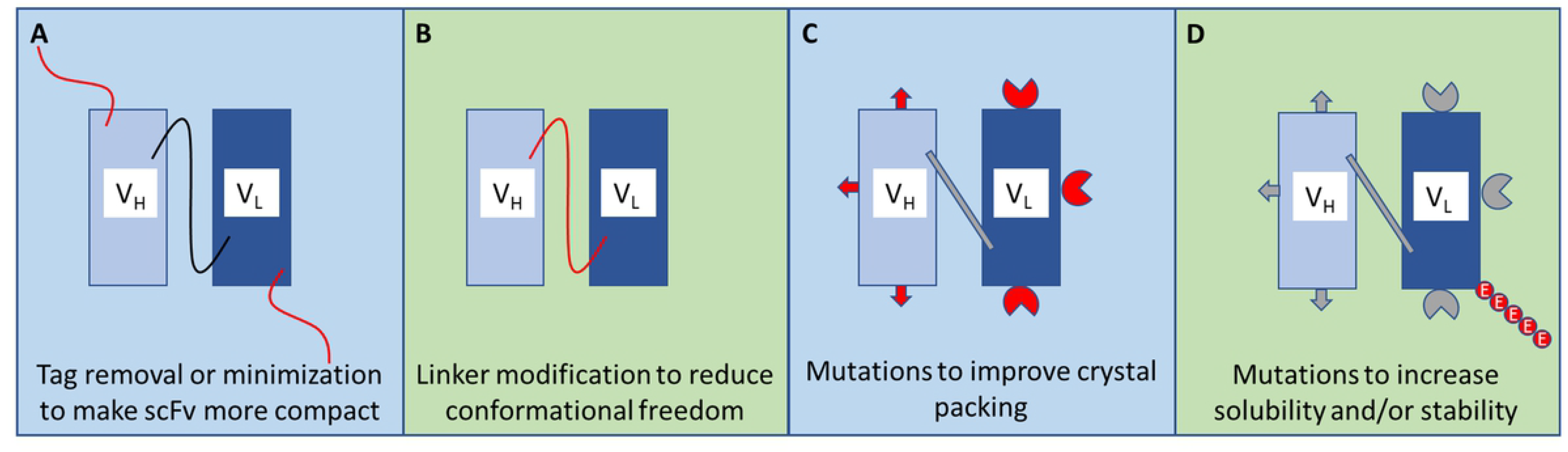
Summary of potential strategies to improve scFv as crystallization chaperones. (A) Trimming of scFv to standard Fv boundaries prior to setting up crystallization trials, since tags used for purification and/or assays are rarely observed in crystal structures. (B) Alternative linkers should be explored for connecting the V_H_ and V_L_ domains, since the widely used flexible glycine serine linkers are rarely observed in crystal structures. (C) Mutations made to surface residues (non-CDR) to promote crystal packing, targeting either surface entropy reduction (38) or introduction of crystal contacts found in other deposited structures (39). (D) Mutations made to improve solubility and/or stability, which may mean adding a penta peptide tag (40) or a more extensive series of mutations such as that implemented in the PROSS approach (41).

## Methods

### SP140 protein expression and purification

The DNA sequence corresponding to amino acids 688-862 of the SP140 protein (UniProtKB – Q13342) was sub-cloned into the expression vector pNIC-Bio2 adding an N-terminal His_6_ tag and a C-terminal Avi tag, creating the SP140A-c055 construct, Supplementary sequence S1. Expression and purification were performed essentially as described (42), see supplementary methods S1 for a detailed protocol.

### Antibody phage selection and ELISA screening of selected scFv

The phage selection procedure was carried out basically as described (43) but the entire process, including the steps of antigen-phage incubation to trypsin elution, was carried out in ordinary 1.5 mL tubes on a rotator with no automation. The number of washing steps was modified and increased with succeeding selection rounds; six in round one and 12 in round four. Also, the recovered phages were propagated in XL1-Blue *E. coli* between the selection rounds. Re-cloning of the selected material in pool followed by transformation into TOP10 *E. coli*, small-scale expression of 94 randomly picked scFv and subsequent ELISA and sequencing experiments were performed analogues to previously reported (43). The affinity was measured using SPR, Supplementary Method S3 and Figure S1.

### scFv protein expression and purification

The scFv gene was sub-cloned into pNIC-MBP2-LIC, adding an N-terminal TEV cleavable MBP-His_6_ tag, creating the YYSP140A-c009 construct (Supplementary sequence S2). This plasmid was used along with pGro7 (Takara) to transform Shuffle T7 Express cells (NEB). Expression and purification was done using a similar method to that described previously (1, 2), see Supplementary methods S2 and Supplementary Figure S2 for further details.

### Complex isolation

To isolate the scFv:SP140 complex, 3 mg of SP140 were mixed with 1 mg of scFv. The mixture was then injected onto a 104 mL Yarra SEC-2000 column (Phenomenex) connected to a NGC system (Bio-Rad) using 10 mM HEPES, 500 mM sodium chloride, 5 % glycerol, 0.5 mM TCEP, pH 7.5 as the mobile phase, (Figure 1B and supplementary figures S2, S3 and S4). Fractions corresponding to the complex were pooled and concentrated to 7.7 mg /mL.

### Crystallization, data collection and structure determination

Crystallization trials were setup at 20 °C in JCSG-plus (Molecular Dimensions) and Crystal Screen HT (Hampton Research) in Swissci 3 well sitting drop plates (Molecular Dimensions). Crystals appeared in several conditions over 4-7 days. Crystals were mounted in MiTeGen loops and flash frozen in liquid nitrogen using 30 % ethylene glycol as cryoprotectant. After screening the best diffracting crystals were found in a condition containing 1.2 M sodium malonate, 0.5 % jeffamine ED-2003, 0.1 M HEPES pH 7. A data set from a single crystal obtained in this condition was collected at the I24 beamline at the Diamond Light Source. The Xia2 auto-processed images (44) were used to perform molecular replacement using Phaser (45). Further refinement of the structure was done using the Phenix suite of programs (46) in combination with Coot (47) and Molprobity (48), see Table 2. The structure factors and co-ordinates were deposited with the protein data bank, PDB code: 6G8R. Protein structure figures were made using either PyMOL (Schrödinger, LLC.) or CCP4mg (49).

## Acknowledgements

Picaud, S.S. (Cloning SP140A), Pike, A.C.W. (Data Collection at beamline), Pinkas, D.M. (Reviewed structure).

